# Broad and differential animal ACE2 receptor usage by SARS-CoV-2

**DOI:** 10.1101/2020.04.19.048710

**Authors:** Xuesen Zhao, Danying Chen, Robert Szabla, Mei Zheng, Guoli Li, Pengcheng Du, Shuangli Zheng, Xinglin Li, Chuan Song, Rui Li, Ju-Tao Guo, Murray Junop, Hui Zeng, Hanxin Lin

**Author notes:** Corresponding author’s e-mail address: Xuesen Zhao, Hanxin Lin. X.Z and D.C contributed equally to this article. Author order was determined in chronological order of participation in this study.

## Abstract

The COVID-19 pandemic has caused an unprecedented global public health and economy crisis. The origin and emergence of its causal agent, SARS-CoV-2, in the human population remains mysterious, although bat and pangolin were proposed to be the natural reservoirs. Strikingly, comparing to the SARS-CoV-2-like CoVs identified in bats and pangolins, SARS-CoV-2 harbors a polybasic furin cleavage site in its spike (S) glycoprotein. SARS-CoV-2 uses human ACE2 as its receptor to infect cells. Receptor recognition by the S protein is the major determinant of host range, tissue tropism, and pathogenesis of coronaviruses. In an effort to search for the potential intermediate or amplifying animal hosts of SARS-CoV-2, we examined receptor activity of ACE2 from 14 mammal species and found that ACE2 from multiple species can support the infectious entry of lentiviral particles pseudotyped with the wild-type or furin cleavage site deficient S protein of SARS-CoV-2. ACE2 of human/rhesus monkey and rat/mouse exhibited the highest and lowest receptor activity, respectively. Among the remaining species, ACE2 from rabbit and pangolin strongly bound to the S1 subunit of SARS-CoV-2 S protein and efficiently supported the pseudotyped virus infection. These findings have important implications for understanding potential natural reservoirs, zoonotic transmission, human-to-animal transmission, and use of animal models.

**Importance:** SARS-CoV-2 uses human ACE2 as primary receptor for host cell entry. Viral entry mediated by the interaction of ACE2 with spike protein largely determines host range and is the major constraint to interspecies transmission. We examined the receptor activity of 14 ACE2 orthologues and found that wild type and mutant SARS-CoV-2 lacking the furin cleavage site in S protein could utilize ACE2 from a broad range of animal species to enter host cells. These results have important implications in the natural hosts, interspecies transmission, animal models and molecular basis of receptor binding for SARS-CoV-2.

## Introduction

Coronavirus disease 2019 (COVID-19) was first identified in Dec. 2019 in the city of Wuhan, China (1), and has since spread worldwide, causing ∼2.3 million infected and around 160,000 fatalities as of April 18th, 2020 (https://coronavirus.jhu.edu/map.html). These numbers are still growing rapidly. The global COVID-19 pandemic has caused an unprecedented public health and economy crisis.

COVID-19 is caused by a novel coronavirus, Severe Acute Respiratory Syndrome Coronavirus 2 (SARS-CoV-2; initially named as 2019-nCoV) (2, 3). The origin of SARS-CoV-2 and its emergence in the human population remain mysterious. Many of the early cases were linked to the Huanan seafood and wild animal market in Wuhan city, raising the possibility of zoonotic origin (4). Sequencing analyses showed that the genome of SARS-CoV-2 shares 79.5%, 89.1%, 93.3%, and 96.2% nucleotide sequence identity with that of human SARS-CoV, bat coronavirus (CoV) ZC45, bat CoV RmYN02, and bat CoV RaTG13, respectively, suggesting that SARS-CoV-2 probably has bat origins (2, 3, 5). This finding is not surprising as bats are notorious for serving as the natural reservoir for two other deadly human coronaviruses, SARS-CoV and Middle East respiratory syndrome coronavirus (MERS-CoV), which previously caused global outbreak, respectively (6, 7).

Although SARS-CoV-2 may have originated from bats, bat CoVs are unlikely to jump directly to humans due to the general ecological separation. Other mammal species may have been served as intermediate or amplifying hosts where the progenitor virus acquires critical mutations for efficient zoonotic transmission to human. This has been seen in the emergence of SARS-CoV and MERS-CoV where palm civet and dromedary camel act as the respective intermediate host (7). The Huanan seafood and wild animal market in Wuhan city would otherwise be a unique place to trace any potential animal source; however, soon after the disease outbreak, the market was closed and all the wild animals were cleared, making this task very challenging or even impossible. As an alternative, wide screening of wild animals becomes imperative. Several recent studies identified multiple SARS-COV-2-like CoVs (SL-CoVs) from smuggled Malayan pangolins in China. These pangolin CoVs (PCoV) form two phylogenetic lineages, PCoV-GX and PCoV-GD (8-11). In particular, lineage PCoV-GD was found to carry a nearly identical receptor-binding motif (RBM) in the spike (S) protein to that of SARS-CoV-2 (Fig.1). However, the genome of these pangolin SL-CoVs share only 85.5%-92.4% nucleotide identities with that of SARS-CoV-2. This is in contrast to SARS-CoV and MERS-CoV where CoVs isolated form the intermediate host palm civet and dromedary camel share 99.6% and 99.9% % genome sequence identities with their human counterpart, respectively (12, 13). Therefore, pangolins tested in these studies are not the direct intermediate host for SARS-CoV-2. Whether or not SARS-CoV-2 came from other pangolins or other wild animal species remains to be determined.

**Fig. 1.**
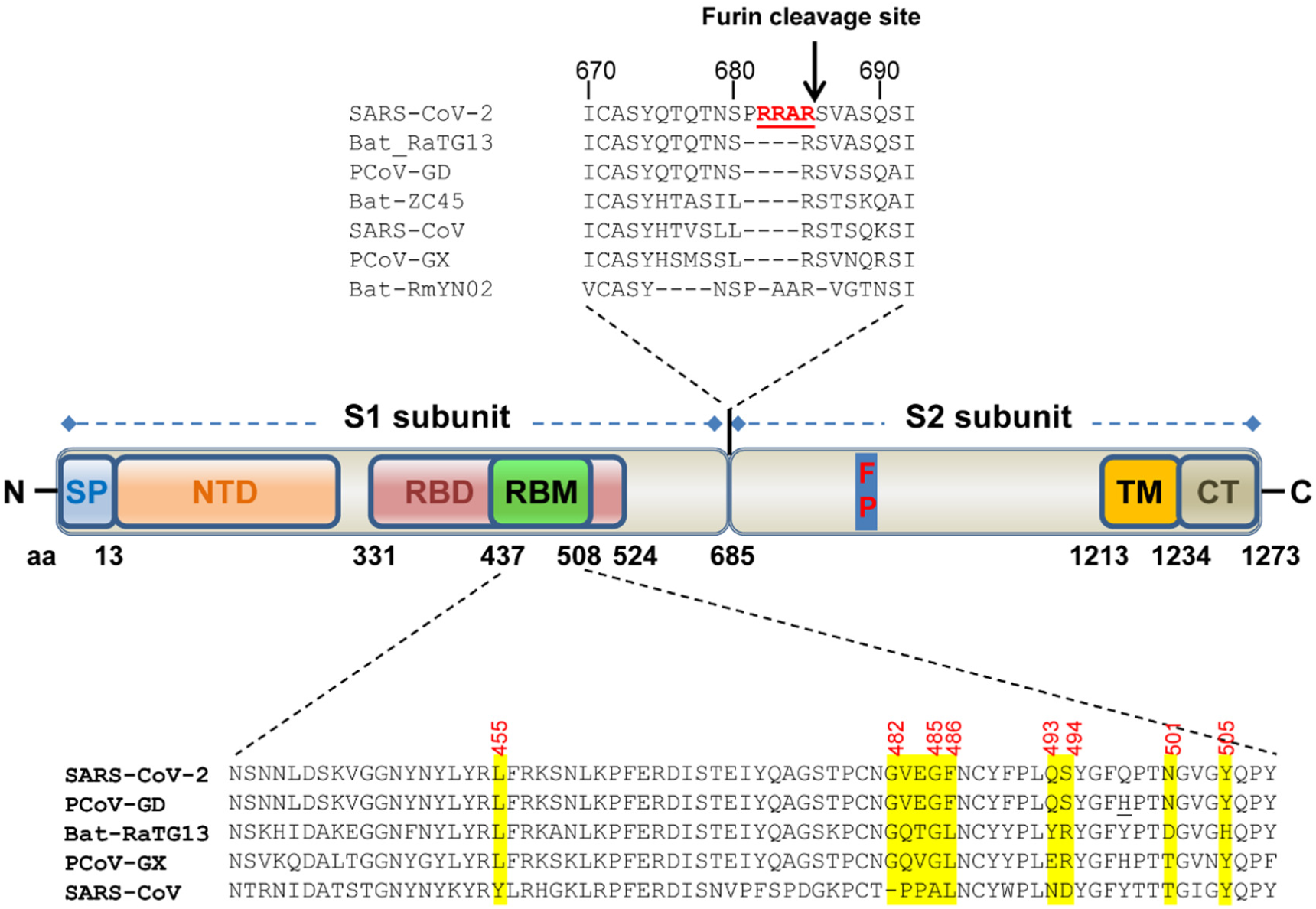
Schematic diagram of domain structures and critical ACE2-binding residues of the spike (S) protein of SARS-CoV-2. SP: signal peptide. NTD: N-terminal domain. RBD: receptor-binding domain. RBM: receptor-binding motif. FP: fusion peptide. TM: transmembrane domain. CT: cytoplasmic tail. The S protein is cleaved into S1 and S2 subunit during biogenesis at the polybasic furin cleavage site (RRAR↓), which is not present in SARS-CoV and other animal SARS-CoV-2-like CoVs. The S1 subunit is required for binding to ACE2 receptor, while the S2 subunit containing a FP mediates membrane fusion. In SARS-CoV-2, the S1 contains NTD and an independently folded domain known as RBD, which harbors a region called receptor binding motif (RBM), that are primarily in contact with receptor. The most critical hACE2-binding residues in the RBM of several SARS-CoV-2-related CoVs are highlighted in yellow and referred from the crystal structure of RBD-hACE2 complex (Shang et al, Nature). PCoV-GX: pangolin CoV isolate GX-PL4. PCoV-GD: pangolin CoV isolate MP789. The only difference in the RBM between PCoV-GD and SARS-CoV-2 is Q498H (underlined). The GenBank No. for these CoVs is: SARS-CoV-2 (isolate Wuhan-Hu-1, MN908947), SARS-CoV (isolate Tor2, NC_004718.3), bat-ZC45 (MG772933.1), bat-RaTG13 (MN996532.1), PCoV-GX (isolate P4L, MT040333.1), PCoV-GD (isolate MP789, MT084071.1).

S protein driven cellular entry, triggered by receptor recognition, is the major determinant of host range, cell, tissue tropism, and pathogenesis of coronaviruses (14). The S protein of SARS-CoV-2 is a type I membrane glycoprotein, which can be cleaved to S1 and S2 subunit during biogenesis at the polybasic furin cleavage site (RRAR) (Fig.1) (15-18). Previous studies have shown that furin cleavage is not essential for coronavirus-cell membrane fusion, but enhances cell-to-cell fusion (19-23), expands coronavirus cell tropism (24), increases the fitness of sequence variant within the quasispecies population of bovine CoV (25). Recent studies indicated that the cleavage at the S1/S2 boundary by furin in virus-producing cells is a critical prime step that facilitates conformation change triggered by receptor binding during virus entry and subsequent fusion-activating cleavage at the S2′site, which is located immediate upstream of fusion peptide in S2 subunit (18, 24, 26). Also, furin cleavage in HA was found to convert avirulent avian influenza virus isolate to a highly pathogenic isolate (27). Interestingly, this cleavage site is not present in the S protein of SARS-CoV, bat SL-CoVs or pangolin SL-CoVs identified so far (5, 15). Besides furin-mediated cleavage in virus-producing cells, SARS-CoV-2 S protein is also cleaved for fusion activation by cell surface protease TMPRSS2 and lysosomal proteases, e.g. cathepsin L, during virus entering target cells (15, 18).

During cell entry, S1 binds to the cellular receptor, subsequently triggering a cascade of events leading to S2-mediated membrane fusion between host cells and coronavirus particles (28). S1 protein contains an independently folded domain called the receptor binding domain (RBD), which harbors an RBM that is primarily involved in contact with receptor (Fig. 1). Human ACE2 (hACE2) has been identified as the cellular receptor for both SARS-CoV-2 (3, 15, 17, 29) and SARS-CoV (30). In addition to hACE2, ACE2 from horseshoe bat (*Rhinolophus alcyone*) was found to support cell entry of SARS-CoV-2 S-mediated VSV-based pseudotyped virus (15). By using infectious virus it has also been shown that ACE2 from Chinese horseshoe bat (*Rhinolophus sinicus*), civet and swine, but not mouse, could serve as functional receptors (3). However, in this infection system, the entry step was coupled with other steps during virus life cycle, i.e. viral genome replication, translation, virion assembly and budding, and thus the receptor activity of these animal ACE2 orthologs were not directly investigated.

In an effort to search for potential animal hosts, we examined the receptor activity of ACE2 from 14 mammal species, including human, rhesus monkey, Chinese horseshoe bat (Rs bat), Mexican free-tailed bat (Tb bat), rat, mouse, palm civet, raccoon dog, ferret badger, hog badger, canine, feline, rabbit, and pangolin for SARS-CoV-2 and a mutant virus lacking the furin cleavage site in the S protein. Our results show that multiple animal ACE2 could serve as receptors for SARS-CoV-2 and the SARS-CoV-2 mutant. ACE2 of human/rhesus monkey and rat/mouse exhibited the highest and lowest receptor activity, respectively, with the other 10 ACE2s exhibiting intermediate activity. The implications of our findings were discussed in terms of the natural reservoir, zoonotic transmission, human-to-animal transmission, animal health, and animal model.

## Results

### Human ACE2 serves as a functional receptor for SARS-CoV-2

To examine the receptor activity of human ACE2 (hACE2) for SARS-CoV-2, we first established a HIV-based pseudotyped virus entry system. This system has been widely used in studies of coronavirus entry. To improve the expression level of S protein and the yield of pseudotyped virus, a codon-optimized S gene based on the sequence of isolate Wuhan-Hu-1 (2) was synthesized and used for production of pseudotyped virus as previously described for other human coronaviruses (HCoVs), including SARS-CoV, MERS-CoV, NL63, 229E, and OC43 (31, 32). The pseudotyped virus was then used to infect 293T cells transfected with either empty vector, or a plasmid expressing APN (receptor for HCoV-229E), DDP4 (receptor for MERS-CoV), ACE1 or hACE2. Two days post-infection, the luciferase activity was measured. As shown in Fig. 2A, only hACE2 was able to efficiently support virus entry. The entry of SARS-CoV-2, but not influenza virus A (IVA) or HCoV-43, was blocked by antibody against hACE2 in a dose-dependent manner (Fig. 2B). We also performed a syncytia formation assay to assess the membrane fusion triggered by hACE2-S binding. As shown in Fig. 1C, syncytia formation was only seen for cells expressing hACE2, but not hACE1, mixed with cells expressing the S protein of SARS-CoV-2 or SARS-CoV. These results confirm that hACE2 is the *bone fide* entry receptor for SARS-CoV-2.

**Fig. 2.**
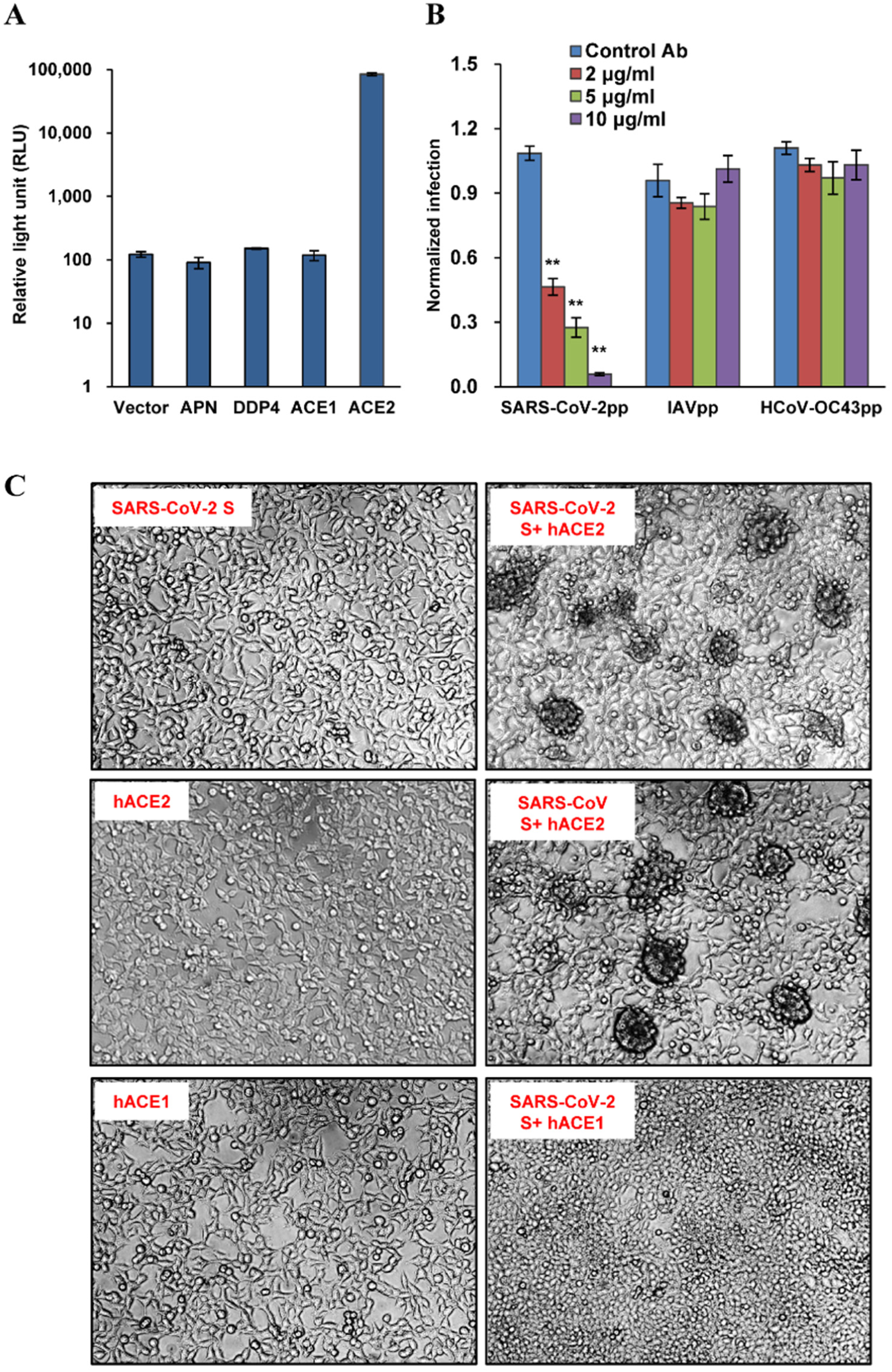
Human ACE2 served as receptor for SARS-CoV-2. (A) ACE2 supported HIV-Luc-based pseudotyped virus entry. 293T cells were transfected with empty vector pcDNA3.1, APN (receptor for HCoV-229E), DDP4 (receptor for MERS-CoV), ACE1 or ACE2. At 48 h post transfection, the cells were infected by SARS-CoV-2 S protein pseudotyped virus (SARS-CoV-2pp). At 48 h post infection, luciferase activity was measured. (B) Human ACE2 antibody inhibited virus entry at a dose-dependent manner. 293T cells were transfected with ACE2. At 48 h post transfection, the cells were pre-incubated with indicated concentration of hACE2 antibody or control antibody (anti-IDE) for 1 h, and then infected by pseudotyped virus of SARS-CoV-2, Influenza virus A (IAVpp) or human coronavirus (HCoV) OC43 (HCoV-OC43pp) in the presence of indicated concentration of hACE2 antibody or control antibody (anti-IDE) for another 3 h, then the virus and antibodies were removed. At 48 h post infection, luciferase activity was measured and normalized to the control antibody for SARS-CoV-2pp. Error bars reveal the standard deviation of the means from four biological repeats. (C) Syncytia formation assay. 293T cells transfected with a plasmid the expressing S protein of SARS-CoV-2 or SARS-CoV were mixed at a 1:1 ratio with those cells transfected with a plasmid expressing ACE1 or ACE2. Twenty-four hours later, syncytia formation was recorded.

### Multiple animal ACE2 orthologs serve as receptors for SARS-CoV-2 and SARS-CoV-2 mutant with S protein lacking the furin cleavage site

To test if other animal ACE2 orthologs can also be used as receptor for SARS-CoV-2, we cloned or synthesized ACE2 from rhesus monkey, Chinese horseshoe bat (Rs bat), Mexican free-tailed bat (Tb bat), rat, mouse, palm civet, raccoon dog, ferret badger, hog badger, canine, feline, rabbit, and pangolin. These animals were chosen as being either the proposed natural hosts for SARS-CoV-2 (bat, pangolin) (3, 10), intermediate hosts for SARS-CoV (civet, raccoon) (12), common animal model (rat, mouse, monkey), or household pets (canine, feline, rabbit). These ACE2 molecules were transiently expressed in 293T cells (Fig.3A), which were then infected with pseudotyped virus of SARS-CoV-2 (SARS-CoV-2pp). The luciferase activity was measured and normalized to hACE2 (Fig. 3B). The results showed that (1) ACE2 of human and rhesus monkey were the most efficient receptors; (2) ACE2 of rat and mouse barely supported virus entry (<10% of hACE2); (3) the receptor activities of the other 10 animal ACE2s were between human/monkey and rat/mouse. Among these, ACE2 of canine, feline, rabbit and pangolin could support virus entry at levels >50% of hACE2.

**Fig. 3.**
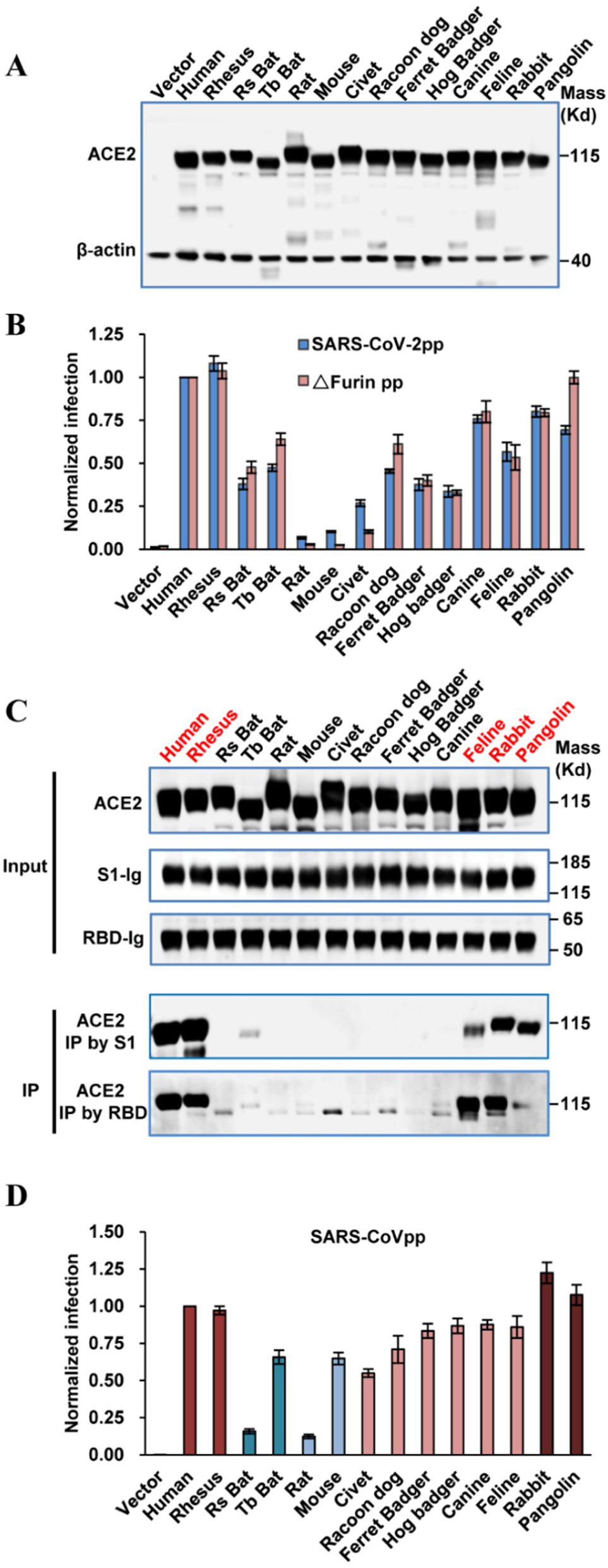
Multiple ACE2 orthologues served as receptors for SARS-CoV-2. (A) Transient expression of ACE2 orthologs in 293T cells. The cell lysates were detected by western blot assay, using an anti-C9 monoclonal antibody. (B) HIV-Luc-based pseudotyped virus entry. 293T cells were transfected with ACE2s orthologs. At 48 h post transfection, the cells were infected by the pseudotyped virus of wildtype SARS-CoV-2 or mutant ΔFurin. At 48 h post infection, luciferase activity was measured and normalized to human ACE2, respectively. Error bars reveal the standard deviation of the means from four biological repeats. (C) IP assay. The upper panel showed the input of ACE2 protein with C9 tag, S1 and RBD with IgG tag. The lower panel showed the ACE2 pulled down by S1-Ig or RBD-Ig fusion protein. (D) SARS-CoV spike-mediated entry. 293T cells were transfected with ACE2s orthologs. At 48 h post transfection, the cells were infected by the pseudotyped virus of SARS-CoV. At 48 h post infection, luciferase activity was measured and normalized to human ACE2, respectively. Error bars reveal the standard deviation of the means from four biological repeats.

To examine receptor binding ability, we performed immunoprecipitation (IP) analysis by using both S1 and receptor binding domain (RBD) as probe. Among the 14 different ACE2s tested, ACE2 from human, monkey, feline, rabbit and pangolin exhibited significant and consistent association with S1 and RBD (Fig.3C). Importantly, these ACE2s correspond to the group of ACE2s that supported the most efficient virus entry (Fig.3B). The Lack of significant entry reduction in 293T cells of furin mutant virus was likely due to the redundancy of cellular proteases, e.g. endosomal cathepsin, that promote membrane fusion in endosome. It has been proposed that MERS-CoV mutant having uncleaved S proteins enter cells via late endosome/lysosome (24). Two recent studies confirmed that furin cleavage of SARS-CoV-2 S protein was required for efficient entry into human lung cells (18, 33).

A striking difference between SARS-CoV-2 and animal SL-CoVs is the presence of a polybasic furin cleavage site at the S1/S2 boundary of the S protein (Fig.1). Here, we generated a SARS-CoV-2 S gene mutant with the furin cleavage site deleted to mimic the bat SL-CoV CZ45. This S mutant has been previously demonstrated to express a full-length non-cleaved S protein during biogenesis in cells (17). Pseudotyped virus with this mutant S protein was produced and used to infect ACE2-transfected 293T cells. Similar or slightly higher efficiency were observed for the mutant S protein-mediated pseudoviral infection in cells transfected with all the animal ACE2s, except for mouse, rat and civet where the mutant S protein mediated a slight lower efficiency of infection. Interesting, pangolin ACE2 was now as efficient as hACE2 for supporting mutant virus entry (Fig.3B).

We also tested the receptor usage of these 14 ACE2 by SARS-CoV (Fig.3D). The results indicated that ACE2 of Rs bat and rat were the poorest receptors (<20% of hACE2), while the other ACE2s could support SARS-CoV entry at levels >50% of hACE2. Interestingly, ACE2 of rabbit and pangolin were even more efficient than hACE2 for supporting SARS-CoV entry. Together, these results demonstrated that SARS-CoV-2 and its mutant virus lacking furin cleavage site, as well as SARS-CoV, could use multiple animal ACE2s as receptor.

### Molecular basis of different ACE2 receptor activities

To help understand the molecular basis of different ACE2 receptor activities, we first examined the overall sequence variation between these ACE2s. For this purpose, we constructed a phylogenetic tree based on the nucleotide sequences of ACE2s (Fig.4). Interestingly, the phylogenetic clustering of ACE2s is correlated with their abilities to support SARS-CoV-2 entry. For example, ACE2s in subclade IIA (human, rhesus monkey and rabbit) and IIB (rat and mouse) were the most efficient and poorest receptor, respectively, while ACE2s in clade I (from the remaining animals) were intermediate between subclades IIA and IIB. This correlation suggests that sequence variations that define for speciation are responsible for observed differences in receptor activity.

**Fig. 4.**
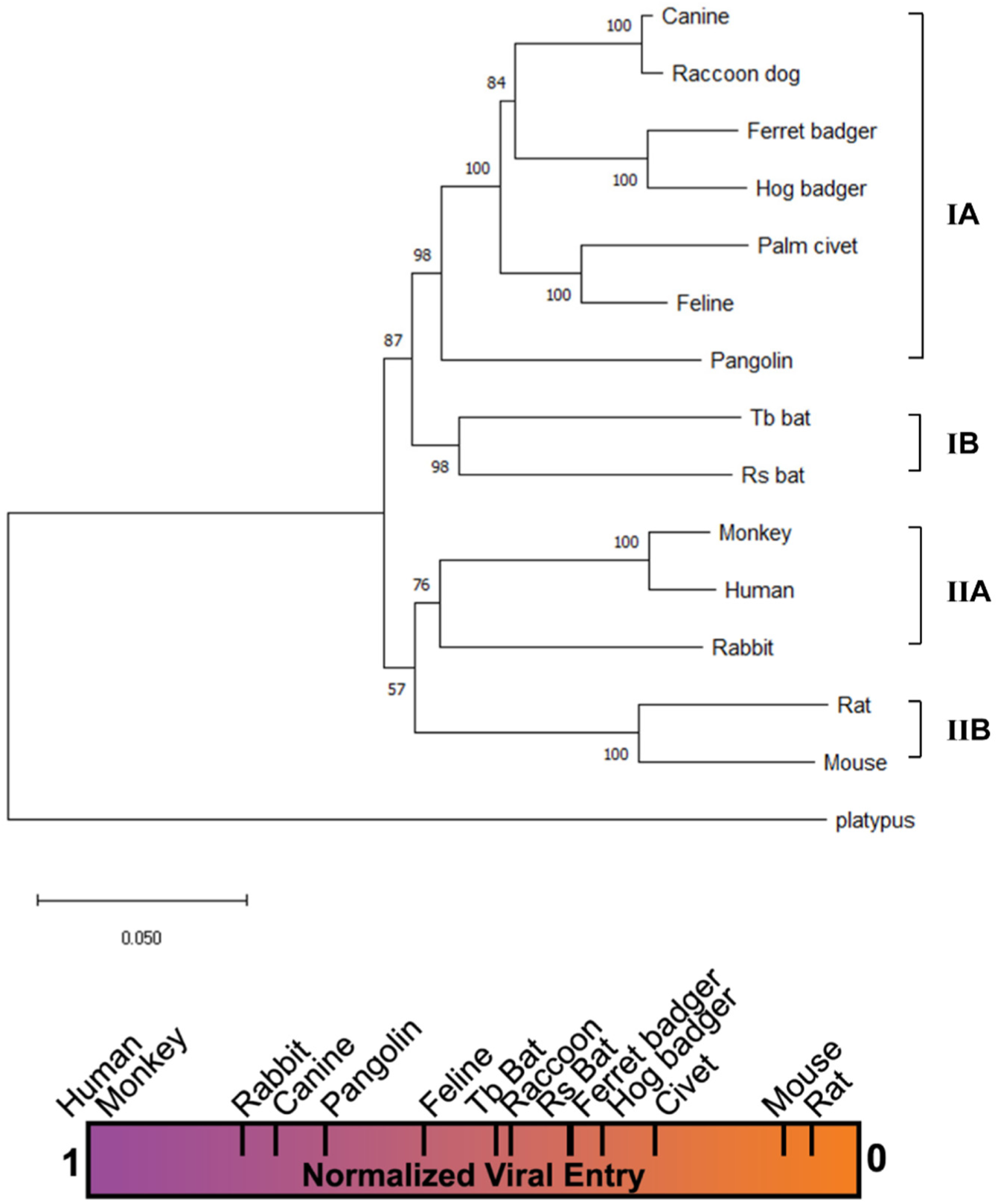
Phylogenetic clustering of ACE2s correlates with their receptor activities. **Upper panel**: phylogram tree of 14 ACE2s. The tree was constructed based on nucleotide sequences using the Neighbor-joining method implemented in program MEGA X. The percentage of replicate trees in which the associated taxa clustered together in the bootstrap test (1000 replicates) are shown next to the branches. The tree was rooted by ACE2 of platypus (*Ornithorhynchus anatinus*). The taxonomic orders where these animals are classified are shown on the right-hand side of the tree. **Lower panel**: a heat bar summarizing the relative levels of pseudotyped virus entry supported by different animal ACE2s.

Next, based on the published crystal structures of hACE2-RBD complex we compared amino acid sequences of ACE2 receptors, focusing on 23 critical residues in close contact with RBD of SARS-CoV-2 (16, 34, 35) (Fig. 5). Two obvious patterns were observed. First, hACE2 and rhesus monkey ACE2 are identical at all critical residues for RBD interaction. This explains why rhesus monkey ACE2 supported virus entry as efficient as hACE2 (Fig.3B). Second, since rat and mouse merely support virus entry, the three substitutions (D30N, Y83F and K353H) that are only seen in rat and mouse ACE2s may be the key.

**Fig. 5.**
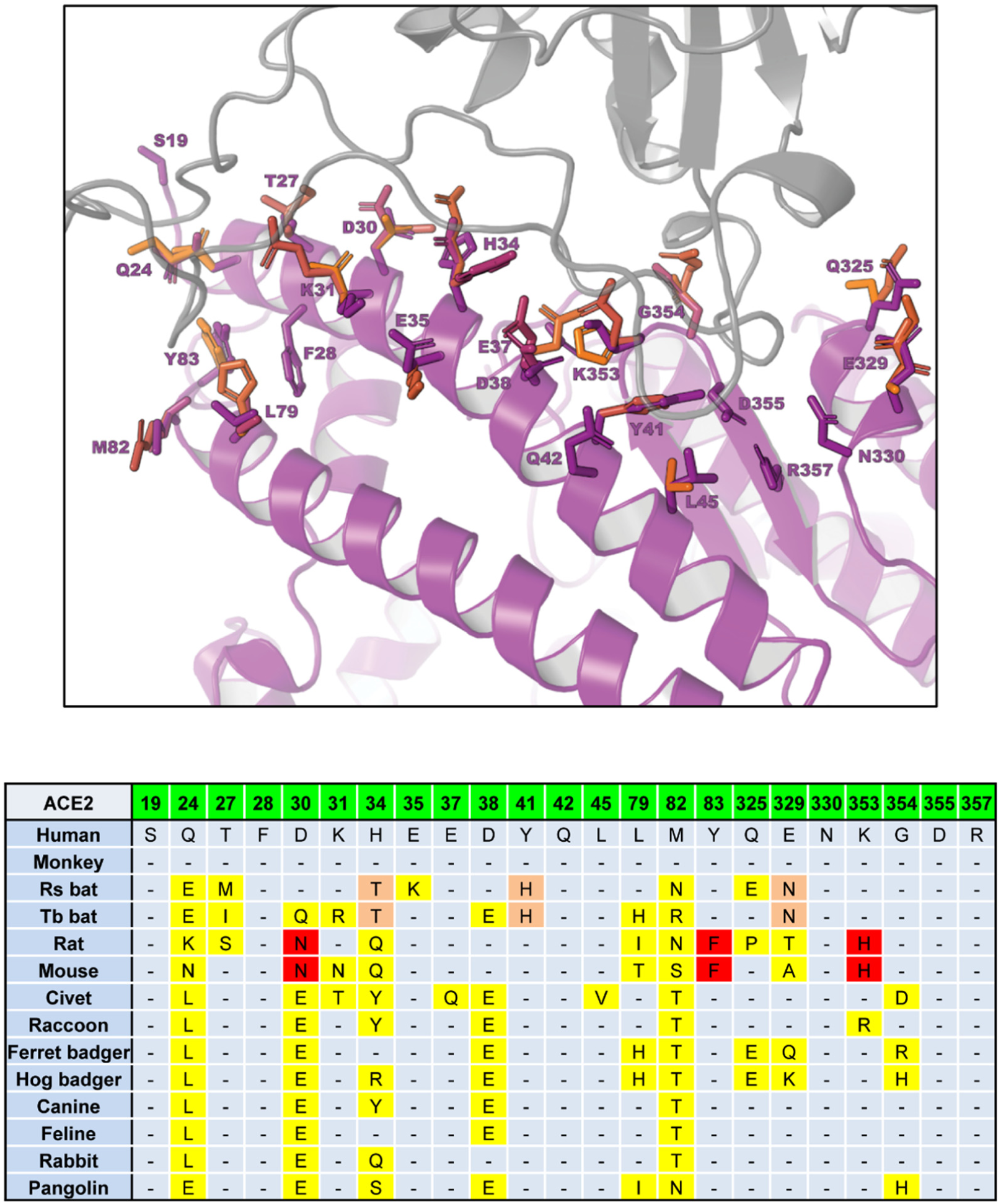
Critical RBD-binding residues in ACE2 orthologs. **Upper panel**: 23 RBD-binding residues at the contact interface between hACE2 and RBD of SARS-CoV-2. Human ACE2 (PDB: 6VW1) in the bound conformation was extracted from the SARS-CoV-2 RBD/ACE2 complex and used as a template for homology modeling (16). **Low panel**: Critical RBD-binding residues in ACE2 orthologs. Residue substitutions highlighted in red and orange are those unique to both mouse and rat ACE2s and both bats, respectively. The rest of residue substitutions are highlighted in yellow.

To further explain the different receptor activities, we used homology-based structure modeling to analyze the effect of residue substitutions at the atomic level. Structure models of 14 ACE2s were generated based on the crystal structure of SARS-CoV-2 RBD/ACE2 complex (16). The effects of critical residue substitutions were analyzed and are summarized in Table 1. Overall, the predicted effects of residue substitutions in ACE2s were consistent with corresponding receptor activities. ACE2 of rodents and bats are presented as examples of this analysis (Fig.6).

**Table 1.**
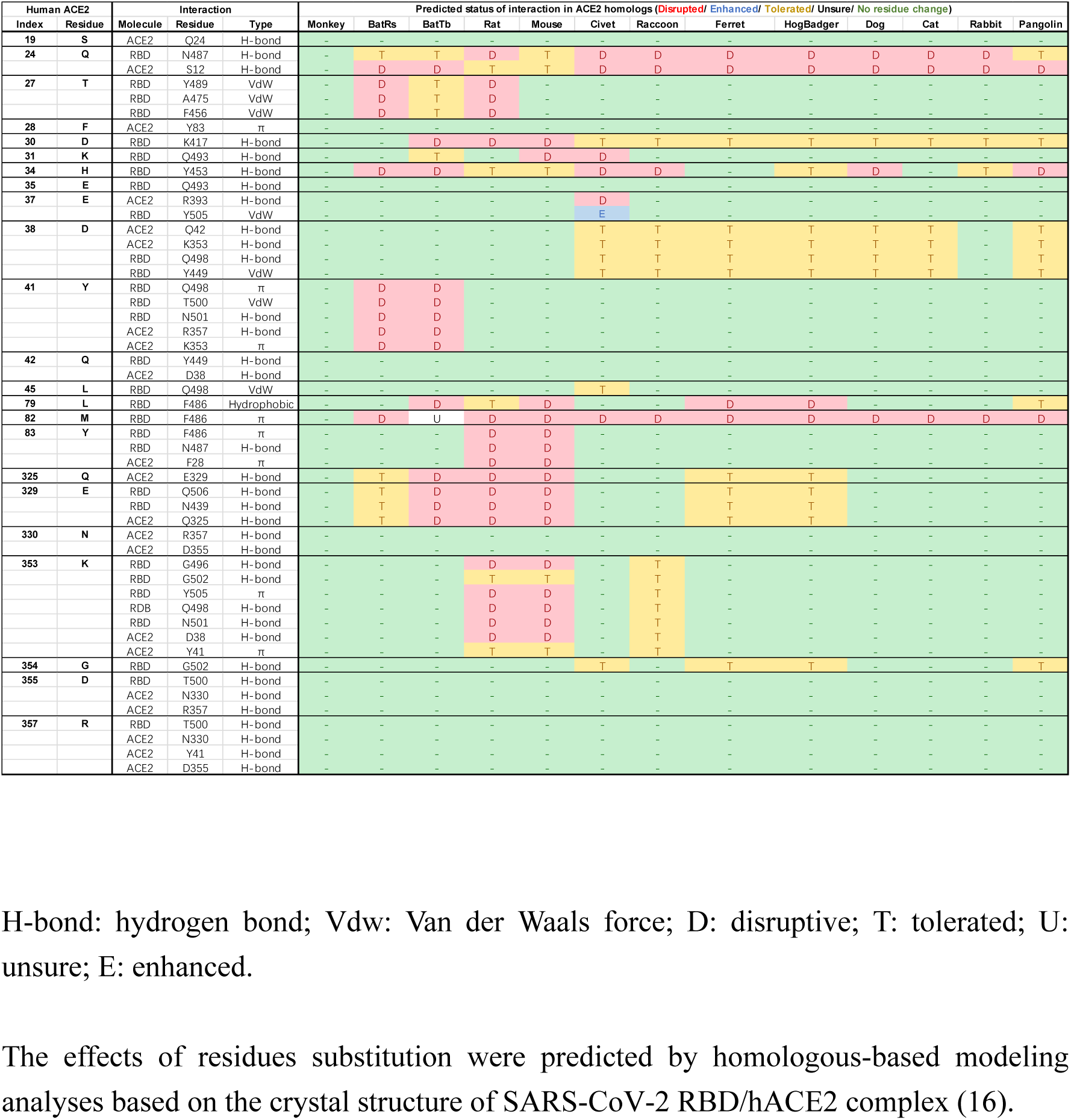
Predicted effect of critical residue substitutions in ACE2 orthologs on the interaction with RBD of SARS-CoV-2

**Fig. 6.**
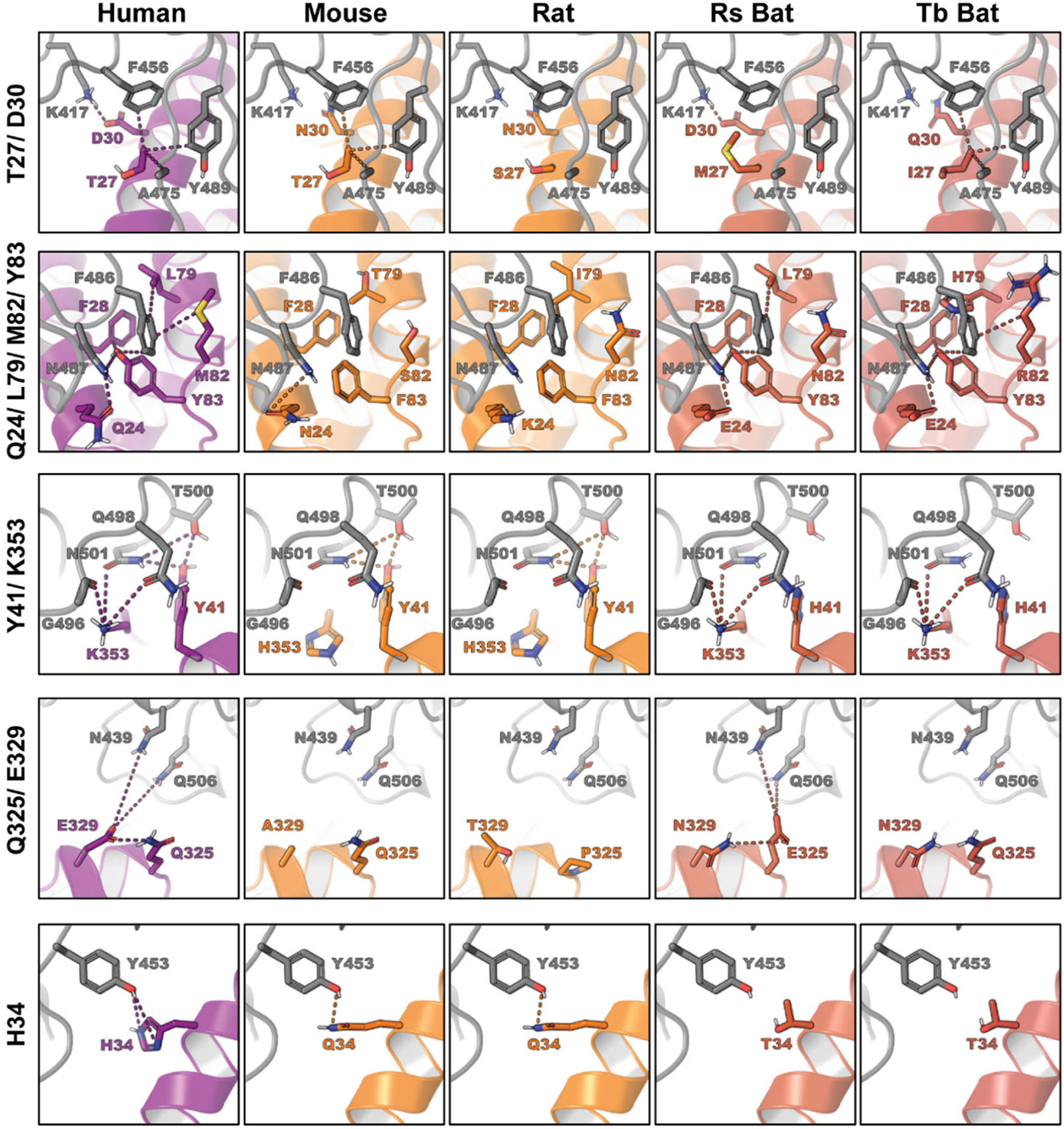
Structural models of key residue substitutions in ACE2 of mouse, rat and bats. Human ACE2 (PDB: 6VW1) in the bound conformation was extracted from the SARS-CoV-2 RBD/ACE2 complex and used as a template for homology modeling (16). ACE2 Homology models were generated using the one-to-one threading algorithm of Phyre2 (63). The models were then aligned and compared to the intact SARS-CoV-2 RBD/ ACE2 complex in PyMOL.

First, we examined the rodent-unique substitutions D30N, Y83F and K353H as they may play a key role in rat and mouse ACE2 inactivity. In humans the residues at all three of these positions directly contact the RBD via hydrogen bonds. D30 contacts K417, Y83 contacts N487, and K353 appears to be at the center of a hydrogen bond network spanning seven RBD residues (Y449, G496, Q498, T500, N501, G502, Y505) and eight ACE2 residues (D38, Y41, Q42, N330, K353, G354, D355, R357). The D30N, Y83F and K353H substitutions are all predicted to disrupt these interactions in rat and mouse ACE2 (Figure 6). This is consistent with previous reports which pinpoint K353 as an important hotspot for both SARS-CoV-2 (16) and SARS-CoV (36) binding. It has been experimentally demonstrated that introduction of K353H into hACE2 significantly reduces binding to SARS-CoV S1; in contrast, introduction of H353K into rat ACE2 significantly increases binding to SARS-CoV S1 (37). Our homology models indicate that other residue substitutions may also be contributing to the low viral entry activity in mouse and rat ACE2. Substitutions Q24N, Q27S, M82N, Q325P and E329T in rat ACE2, and L79T, M82S and E329A in mouse ACE2, are all predicted to disrupt interactions with RBD residues (Fig. 6 and Table 1).

Both Bat ACE2s are also inefficient receptors for viral entry (Fig.3B). Since the profile of residues at the receptor/RBD interface is significantly different from rat and mouse ACE2, we examined other bat-specific residue substitutions that may be contributing to receptor dysfunction. There are 8 and 10 critical residue substitutions in the Rs bat and Tb bat ACE2s, respectively (Fig.5). Among these, we examined the substitutions at positions Y41, H34 and E329 as they are only seen in bat ACE2s. The Y41H substitution in both bat ACE2s appears to be disrupting the same H-bond network that was disrupted by K353H in rat and mouse ACE2. Although Y41 is not as centrally located in the H-bond network as K353, it directly contacts N501 from the RBD, which is the same residue that is stabilized by K353. A second interaction which appears to be disrupted in only bat ACE2s occurs at position H34. In humans, H34 forms a H-bond with Y453 from the RBD, which is broken through a H34T substitution in bat ACE2s. Finally, the bat-unique substitution E329N appears to be disrupting H-bonds connecting two ACE2 residues (E329, Q325) and two RBD residues (N439, Q506). In Tb bat ACE2, all connections in the H-bond network are disrupted by the single E329N substitution, however the H-bond network is predicted to be restored by an additional substitution, Q325E in Rs bat. In addition, other residue substitutions, i.e. T27M and M82N in Rs bat, and D30Q and L79H in Tb bat, are also disruptive (Fig. 6).

These results reveal that the poor and low receptor activity of rodent ACEs and bat ACE2 are resulted from a broken interaction network by a key residue substitution, i.e. K353H in rodents and Y41H in bats, and additive disruptive effects by multiple residue substitutions.

## Discussion

In this study, we examined the receptor activity of 14 ACE2 orthologues. The results suggested that wild type and mutant SARS-CoV-2 lacking the furin cleavage site in S protein could use ACE2 from a broad range of animal species to enter host cells. Below we discuss the implication of our findings in terms of natural reservoir, zoonotic transmission, human-to-animal transmission, animal health, and animal model.

### Implication in natural reservoirs and zoonotic transmission

Among the 14 ACE2s tested here, hACE2 and rhesus monkey ACE2 are the most efficient receptors, suggesting that SARS-CoV-2 has already been well adapted to humans. In addition, ACE2s of other animals, except mouse and rat, could also support SARS-CoV-2 entry (Fig.3B). Although these data were obtained by using HIV1-based pseudotyped virus, for ACE2 of Rs bat, civet, and mouse, the data is consistent with *in vitro* infection data using infectious virus (3). Receptor usage by coronaviruses has been well known to be a major determinant of host range, tissue tropism, and pathogenesis (14, 38, 39). It is therefore reasonable to assume that SARS-CoV-2 would be able to infect all these animals. As a matter of fact, several *in vivo* infection and seroconversion studies have confirmed that SARS-CoV-2 can infect rhesus monkey (40), feline, ferret, and canine (41, 42). Our findings are also in line with the concordance between ACE2 receptor usage by SARS-CoV pseudotyped virus and susceptibility of the animals to SARS-CoV infection. As shown in Fig.3D, ACE2 of rhesus monkey, mouse, civet, ferret badger, raccoon and feline could support SARS-CoV pseudotyped virus entry; concordantly, all these animals are susceptible to native SARS-CoV virus infection (12, 43-46).

Among all those wild animals that are potentially infected by SARS-CoV-2, bat and pangolin have already been proposed to be the natural reservoirs as closely related SL-CoVs have been identified in bats (2, 3, 5) and pangolins (8-11). A recent study has shown that bat SL-CoV RaTG13 could use hACE2 as receptor, consistent with the presence of several favorable hACE2-binding residues (aa 455, 482-486) in the receptor binding motif (RBM) of the S protein (Fig.1) (16). For pangolin SL-CoVs, lineage PCoV-GD has only one non-critical amino acid substitution (Q483H) in the RBM when compared to SARS-CoV-2 (Fig.1) (10). Therefore, PCoV-GD most likely can also use hACE2 and other animal ACE2s as functional receptors.

We also tested the receptor usage by a SARS-CoV-2 mutant that lacks the furin cleavage site at the S1/S2 boundary. Our result showed that the mutant virus behaved similarly to the wt virus. Namely, the entry of mutant virus could also be supported by those animal ACE2s that supported the entry of wt virus. This result is similar to another study that used the S gene mutant but in a MLV-based pseudotyped virus system (17) and the role of furin cleavage during coronavirus infection. Furin cleavage is not essential for coronavirus-cell membrane fusion, but enhances cell-to-cell fusion (19-22, 47). This could provide certain level of advantage during infection. For example, in the quasispecies population of bovine CoV, a minor sequence variant with a polybasic furin-like cleavage site in the S2 subunit quickly dominated the population even after a single passage in cells (25). However, by using pseudotyped virus system, which is a single cycle infection system, we may not see the advantage. Still, ours result unequivocally showed that SARS-CoV-2 without this cleavage site could use multiple animal ACE2s as receptors to enter cells. As there is a need to continuously search for potential intermediate hosts for SARS-CoV, results presented here can help significantly narrow down the scope of potential targets.

Collectively, our results highlight the potential of these wild animals to serve as natural reservoirs or intermediate hosts for SARS-CoV-2 and its progenitor, the risk of zoonotic transmission of animal SL-CoVs to human, and the necessity of virus surveillance in wild animals.

### Implication in human-to-animal transmission and animal health

Among those animal species tested here, canine and feline are of special concern as they are often raised as companion pets. Our data indicate that ACE2 of canine and feline could support SARS-CoV-2 pseudotyped virus entry quite efficiently (>50% of hACE2, Fig.3B), raising the alarming possibility of virus transmission from infected human to these pets or potentially vice versa. As a matter of fact, there was a recent report that a Pomeranian dog in Hong Kong tested weakly positive for SARS-CoV-2 while maintaining an asymptomatic state. The genome of the virus isolated from this dog has only three nucleotide changes compared to the virus isolated from two infected persons living in the same household, suggesting that this dog probably acquired the virus from the infected owners (48). Our results are further supported by two additional studies. One study showed that both dog and cat were susceptible to SARS-CoV-2 infection. While the virus replicated poorly in dogs, it replicated efficiently in cats and was able to transmit to unaffected cats that were housed with them (41). The other study revealed that 14.7% of cat sera samples collected in Wuhan city after the outbreak were positive for antibody against SARS-CoV-2, demonstrating that many cats were infected during the outbreak, most likely from infected humans in close contact (42). Domestic cats are also susceptible to SARS-CoV infection (43) and human-to-cat transmission was evident during the SARS-CoV outbreak in 2003 in Hong Kong (49). These findings were also in agreement with our results that ACE2 of cat and dog could serve as receptor for SARS-CoV (Fig.3D).

As described above, it seems that dogs are not as susceptible as cats to SARS-CoV-2 (41, 48). Interestingly, this is in agreement with results from IP analysis that showed cat ACE2 could bind to S1 or RBD more efficiently than dog ACE2 (Fig.3C). Structural models further suggest that, at those critical RBD-binding residues, dog and cat ACE2 share 4 substitutions (Q24L, D30E, D38E, and M82T), while dog ACE2 has an additional substitution, H34Y (Fig.5). Based on structural modeling, both Q24L and M82T are predicted to be disruptive, while both D30E and D38E are tolerable (Table 1). H34Y in dog ACE2 is predicted to disrupt the hydrogen bond with Y453 of RBD (Table 1). These atomic interactions explain why dog ACE2 binds to S1 or RBD less efficiently compared to cat ACE2, and both are less efficient than human ACE2.

In addition to cat and dog, rabbits are also often raised as household pets. Our results indicate that rabbit ACE2 is an efficient receptor (Fig. 3B and 3C), suggesting that rabbit may be more susceptible to SARS-CoV-2 infection than cat.

Currently, there is no evidence that infected pets can transmit the virus back to human; however, this may be possible and should be investigated. Out of an abundance of caution it would be best when one is infected to have both human and pets quarantined, and the pets tested as well.

### Implication in animal model

Animal models are essential for study of pathogenesis, vaccinology and therapeutics of viral pathogens. Rodents are probably the most common and amenable animal models because of low cost, easy handling, defined genetics, and the possibility of scalability (50). However, our results showed that both mouse and rat ACE2 are poor receptors for SARS-CoV-2 (Fig.3B and 3C), suggesting that they are probably resistant to infection. Actually, this has been verified by using infectious SARS-CoV-2 to infect mouse ACE2-transfected cells (3) or mice (51). Genetically engineered mice expressing hACE2 were previously developed as an animal model for SARS-CoV (52). This model has been tested recently for SARS-CoV-2, and found to be susceptible to SARS-CoV-2 infection and development of interstitial pneumonia (51), a common clinical feature of COVID-19 patients (53). Human ACE2-transgenic mice therefore represent useful animal models. However, because of the high demand, and discontinuance due to the disappearance of SARS-CoV in the human population after 2004, it is expected that this mouse model will be in short supply (54). Alternative methods should be sought to develop a mouse-adapted SARS-CoV-2 strain. Mouse-adapted SARS-CoV strains were developed by serial passage of virus in mice (55, 56). However, this method may not work for SARS-CoV-2 as mouse ACE2 still supports some entry for SARS-CoV (Fig.3D), but not SARS-CoV-2. An alternative way to make a mouse-adapted SARS-CoV-2 strain could be achieved by rational design of the S gene. Based on the structural model, we know that receptor dysfunction of mouse ACE2 is due to disruptive D30N, L79T, M82S, Y83F, E329A and K353H substitutions (Fig.5, Fig.6 and Table 1). Therefore, by specifically introducing mutations into the RBM of S gene it may be possible to restore or at least partly restore interactions with these ACE2 substitutions. Consequently, the engineered virus may be able to efficiently infect wildtype mice.

To date, several animals (i.e. rhesus monkey, ferret, dog, cat, pig, chicken and duck) have been examined as potential animal models for SARS-CoV-2 (40, 41). Although the rhesus monkey, ferret and cat may seem to be the promising candidates, none of them are perfect in terms of recapitulation of typical clinical features in COVID-19 patients. Therefore, multiple animal models may be needed. Our results indicate that rabbit ACE2 is a more efficient receptors than other animal ACE2s for both SARS-CoV-2 and SARS-CoV (Fig.3). Therefore, it may be worthy assessing rabbit as a useful animal model for further studies.

## Methods

### Cell lines and antibodies

293T cells and Lenti-X 293T cells were cultured in Dulbecco’s modified Eagle’s medium (DMEM; Gibco) (57). All growth medium was supplemented with 10% fetal bovine serum (FBS), 110 mg/L sodium pyruvate, and 4.5 g/L D-glucose. β-actin antibody and C9 antibody were purchased from Sigma (A2228) and SANTA CRUZ (sc-57432), respectively. A polyclonal antibody against human ACE2 and anti-IDE polyclonal antibody were purchased from R&D Systems (catalog No. AF933 and AF2496, respectively).

### Construction of ACE2 plasmids

ACE2 of human (*Homo sapiens*, accession number NM_001371415.1), civet (*Paguma larvata*, accession number AY881174.1), and rat (*Rattus norvegicus*, accession number NM_001012006.1) were cloned into a modified pcDNA3.1-cmyc/C9 vector (Invitrogen) as previously described (37, 58). ACE2 protein expressed from this vector has a c-myc tag at the N-terminus and a C9 tag at the C-terminus. An Age I site was engineered right downstream of the signal peptide sequence (nt. 1-54) of ACE2. ACE2 protein expressed from this vector has a c-myc tag at the N-terminus and a C9 tag at the C-terminus. ACE2 of Chinese ferret badger (*Melogale moschata*, accession number MT663957), raccoon dog (*Nyctereutes procyonoides*, accession number MT663958), Mexican free-tailed bat (*Tadarida brasiliensis*, accession number MT663956), rhesus monkey (*Macaca mulatta*, accession number MT663960), hog badger (*Arctonyx collaris*, accession number MT663962), New Zealand white rabbit (*Oryctolagus cuniculus*, accession number MT663961), domestic cat (*Felis catus*, accession number MT663959) and domestic dog (*Canis lupus familiaris*, accession number MT663955) were cloned into Age I/Kpn I-digested pcDNA3.1-cmyc-C9 vector previously (Hanxin Lin, Ph. D Thesis Dissertation. “Molecular interaction between the spike protein of human coronavirus NL63 and ACE2 receptor” McMaster University, Health Science Library.https://discovery.mcmaster.ca/iii/encore/record/C__Rb2023203__SMolecular%20interaction%20between%20the%20spike%20protein%20of%20human%20coronavirus%20NL63%20and%20ACE2%20receptor%20Lw%3D%3D%20by%20Hanxin%20Lin__Orightresult__U__X4?lang=eng&suite=def). The nucleotide sequence of ACE2 of Chinese horseshoe bat (*Rhinolophus sinicus*, Rs, accession number KC881004.1) and pangolin (*Manis javanica*, accession number XM_017650263.1) were synthesized and cloned into pcDNA3.1-N-myc/C-C9 vector.

### Construction of plasmids expressing S, S1 and RBD of SARS-CoV-2

The nucleotide sequence of SARS-CoV-2 S gene was retrieved from NCBI database (isolate Wuhan-Hu-1, GenBank No. MN908947). According the method described by Gregory J. Babcock et al (59), the codon-optimized S gene was synthesized, and cloned into pCAGGS vector. The SARS-CoV-2 S gene mutant without the furin cleavage site at the S1/S2 boundary was generated by an overlapping PCR-based method as previously described (60). The S1 subunit (aa 14-685) and RBD (aa 331-524) were cloned into a soluble protein expression vector, pSecTag2/Hygro-Ig vector, which contains human IgG Fc fragment and mouse Ig *k*-chain leader sequence (61). The protein expressed is soluble and has a human IgG-Fc tag.

### Western blot assay

As previously described, the expression of ACE2-C9, S1-Ig, and RBD-Ig fusion proteins were examined by western-blot (61). Briefly, lysates or culture supernatants of 293T cells transfected with plasmid encoding ACE2 orthologs and S1-Ig or RBD-Ig were collected, boiled for 10 min, and then resolved by 4∼12% SDS-PAGE. A PVDF membrane containing the proteins transferred from SDS-PAGE was blocked with blocking buffer (5% nonfat dry milk in TBS) for 1h at room temperature and probed with primary antibody overnight at 4 °C. The blot was washed three times with washing buffer (0.05% tween-20 in TBS), followed by incubation with secondary antibody for 1h at room temperature. After three-time washes, the proteins bounded with antibodies were imaged with the Li-Cor Odyssey system. (Li-Cor Biotechnology).

### Immunoprecipitation (IP) assay

The association between Ig-fused S1 protein or RBD protein and ACE2 protein with C9 tag was measured by IP according to a previously described method (60). Briefly, HEK293T cells were transfected with plasmid encoding ACE2 with Lipofectamine 2000 (Invitrogen). 48 h post-transfection, the transfected 293T cells were harvested and lysed in PBS buffer containing 0.3 % *n*-decyl-β-D-maltopyranoside (DDM, Anatrace). Cell lysates were incubated with Protein A/G PLUS Agarose (Santa Cruz, sc-2003) together with 4μg of S1-Ig or RBD-Ig. Protein A/G agarose were washed three times in TBS/1% Ttriton-X100, resolved by SDS–PAGE, and detected by western blot using anti-C9 monoclonal antibody.

### Production of pseudotyped virus

Following the standard protocol of calcium phosphate transfection, Lenti-X cells in 10-cm plate were co-transfected by 20μg of HIV-luc and 10μg of CoVs spike gene plasmid. At 48h post-transfection, 15 ml supernatant was collected and passed through a 0.45um pore size PES filter. The purified virus was titrated with Lenti-X p24 Rapid Titer Assay (Takara Bio, Cat. No. 632200). The virus was stored at -80 °C for future use.

### Virus entry assay

Each well of 293T cells in a 96-well plate was transfected with 0.1µg of ACE2 plasmid DNA following the standard protocol of Lipofectamine 2000 (Invitrogen). At 48 hours post-transfection, 150 µl of p24-normalized (10 ng) of pseudotype virus was added into each well and incubated at 37°C for 3 hours. The virus was then removed, and 250 µl of fresh medium was added into each well for further incubation. Two days post-infection, the medium was removed and the cells were lysed with 30 µl/well of 1× cell lysis buffer (Promega) for 15 min, followed by adding 50 µl/well of luciferase substrate (Promega). The firefly luciferase activities were measured by luminometry in a TopCounter (PerkinElmer). For each ACE2, four wells were tested in a single experiment, and at least three repeat experiments were carried out. The luciferase activity was expressed as relative light unit (RLU) and normalized to human ACE2 for plotting.

### Syncytial formation assay

293T cells with approximately 90% confluent on 12-well plate were transfected with 1.6μg of plasmid DNA encoding viral S gene or ACE2. At 24 h post-transfection, 293T cells expressing the S protein were mixed at a 1:1 ratio with 293T cells expressing ACE2 and plated on 12-well plate. Multinucleated syncytia were observed 24 h after the cells were mixed.

### Sequence analysis

Multiple alignments of nucleotide or amino acid sequences of the spike gene of coronaviruses and ACE2 orthologs were performed using Clustal X (62). Phylogenetic tree was constructed based on the nucleotide sequences of animal ACE2 using the neighbor-joining algorithm implemented in MEGA X. The tree is drawn to scale with branch lengths in the same units as those of the evolutionary distances used to infer the phylogenetic tree. Evaluation of statistical confidence in nodes was based on 1000 bootstrap replicates. Branches with <50% bootstrap value were collapsed. Platypus ACE2 (*Ornithorhynchus anatinus*, GenBank No. XM_001515547) was used as an outgroup.

### Homology-based structural modeling

Human ACE2 (PDB: 6VW1) in the bound conformation was extracted from the SARS-CoV-2 RBD/ hACE2 complex and used as a template for homology modeling (16). ACE2 Homology models were generated using the one-to-one threading algorithm of Phyre2 (63). The models were then aligned and compared to the intact SARS-CoV-2 RBD/ ACE2 complex in PyMOL (The PyMOL Molecular Graphics System, Version 2.0 Schrödinger, LLC).

